# Prenatal Arsenite Exposure Alters Maternal Cardiac Remodeling During Late Pregnancy

**DOI:** 10.1101/2023.09.28.559986

**Authors:** Nicole Taube, Raihan Kabir, Obialunanma V. Ebenebe, Haley Garbus, Sarah-Marie Alam El Din, Emily Illingworth, Michael Fitch, Nadan Wang, Mark J. Kohr

**Affiliations:** Department of Environmental Health and Engineering, Johns Hopkins Bloomberg School of Public Health, Baltimore, Maryland, United States; Cardiology Division, Department of Medicine, Johns Hopkins University School of Medicine, Baltimore, Maryland, United States

**Keywords:** Arsenic, Maternal exposure, Cardiotoxicity, Cardiovascular Disease, Pregnancy

## Abstract

Exposure to inorganic arsenic through drinking water is widespread and has been linked to many chronic diseases, including cardiovascular disease. Arsenic exposure has been shown to alter hypertrophic signaling in the adult heart, as well as in-utero offspring development. However, the effect of arsenic on maternal cardiac remodeling during pregnancy has not been studied. As such, there is a need to understand how environmental exposure contributes to adverse pregnancy-related cardiovascular events. This study seeks to understand the impact of trivalent inorganic arsenic exposure during gestation on maternal cardiac remodeling in late pregnancy, as well as offspring outcomes. C57BL/6J mice were exposed to 0 (control), 100 or 1000 µg/L sodium arsenite (NaAsO_2_) beginning at embryonic day (E) 2.5 and continuing through E17.5. Maternal heart function and size were assessed via transthoracic echocardiography, gravimetric measurement, and histology. Transcript levels of hypertrophic markers were probed via qRT-PCR and confirmed by western blot. Offspring outcomes were assessed through echocardiography and gravimetric measurement. We found that exposure to 1000 µg/L iAs abrogated normal physiologic growth of the maternal heart during late pregnancy and reduced transcript levels of estrogen receptor alpha (ERα), progesterone receptor membrane component 1 (Pgrmc1) and progesterone receptor membrane component 2 (Pgrmc2). Both 100 and 1000 µg/L iAs also reduced transcription of protein kinase B (Akt) and atrial natriuretic peptide (ANP). Akt protein expression was also significantly reduced after 1000 µg/L iAs exposure in the maternal heart with no change in activating phosphorylation. This significant abrogation of maternal cardiac hypertrophy suggests that arsenic exposure during pregnancy can potentially contribute to cardiovascular disease. Taken together, our findings further underscore the importance of reducing arsenic exposure during pregnancy and indicate that more research is needed to assess the impact of arsenic and other environmental exposures on the maternal heart and adverse pregnancy events.

## INTRODUCTION

Inorganic arsenic (iAs) is a widespread drinking water contaminant and a World Health Organization top chemical of concern^1^. iAs is naturally found in the Earth’s crust, and activities such as mineral extraction, waste processing, pesticide application, and additives to poultry and swine feed have mobilized iAs into food and drinking water supplies^2^. While also found in air and food, the greatest public health threat stems from the ingestion of iAs-contaminated groundwater, and food prepared with such groundwater^2,3^. There are two forms of iAs, trivalent and pentavalent arsenic, with trivalent arsenic being the most toxic form^4^. iAs has been linked to the etiology of many chronic diseases, including cancer, chronic respiratory disease, and cardiovascular disease (CVD) ^5–13^. iAs has been linked to many forms of CVD that include hypertension, atherosclerosis, coronary heart disease, and stroke^14^. Research from our group has demonstrated that chronic iAs exposure induces sex-specific pathological remodeling of the adult heart and alters susceptibility to ischemic injury^15,16^. However, studies have yet to examine the impact of iAs on the cardiovascular system during physiological stressors, such as pregnancy.

During pregnancy, the heart undergoes hypertrophic remodeling, increasing chamber size and wall thickness in response to the increase in blood volume that occurs during pregnancy^17,18^. Blood volume in the pregnant mother increases significantly starting at 6 weeks of gestation, resulting in a 45% increase in blood volume compared to nonpregnant women^19–21^. Cardiac output also increases during pregnancy and remains constant after the end of the second trimester^22,23^. This pregnancy-induced hypertrophy is reversible^24,25^ and regulated by various phases of hormonal signaling during pregnancy. Both progesterone and estradiol increase throughout pregnancy, and they have been shown to modulate cardiac hypertrophy^26,27^. Progesterone is known to induce cardiomyocyte hypertrophy, while estradiol has a biphasic effect, promoting hypertrophy at low concentrations and inhibiting hypertrophy at higher concentrations ^28,29^.

Emerging evidence suggests that the environment plays a role in cardiovascular-related pregnancy complications. Exposure to air pollution and metals have been associated with reduced conception rates, infertility, and increased pregnancy loss^30–32^, suggesting interference with hormone signaling. However, it is unclear how exposure to environmental factors is associated with peripartum cardiomyopathy and other cardiovascular complications during pregnancy, as the mechanism through which the environment impacts the cardiovascular system during pregnancy has yet to be explored. Metals, such as iAs, have negative effects on the cardiovascular system, which undergoes physiological alterations during pregnancy^8,33–36^. However, it is not clear what pathways or physiologic processes are altered due to environmental exposure during pregnancy, and specifically iAs exposure. In the US, 23.8 maternal deaths occur for every 100,000 live births and CVD is the leading cause of pregnancy-related mortality^37,38^. Women who develop complications during pregnancy, such as preeclampsia, hypertension, or gestational diabetes, are at higher risk for adverse pregnancy outcomes and CVD postpartum^39^.

Our previous studies demonstrated that iAs exposure altered cardiac hypertrophy and hypertrophic signaling in the hearts of adult male mice, and we next sought to examine the impact of iAs exposure on maternal cardiac remodeling during pregnancy^15,16^. Pregnant C57BL6/J mice were exposed to either 0, 100 or 1000 µg/L iAs in drinking water during gestation, and a late pregnancy (E17.5) endpoint was examined. We found that pregnant female mice exposed to 1000 µg/L iAs blunted maternal cardiac hypertrophy, decreased transcriptional levels of estrogen receptor alpha (ERα), progesterone receptor membrane component 1 and 2 (Pgrmc1/2), atrial natriuretic peptide (ANP), and protein kinase B (Akt), and decreased protein expression of Akt in the maternal heart during late pregnancy.

## METHODS

### Animals and Arsenic Exposure

This investigation conforms to the *Guide for the Care and Use of Laboratory Animals* published by the National Institutes of Health (NIH; Publication No. 85-23, Revised 2011) and was approved by the Institutional Animal Care and Use Committee of Johns Hopkins University. Timed pregnant female C57BL/6 mice were purchased from Jackson Laboratory (Bar Harbor, ME). Mice (2 per cage) were housed under specific pathogen-free conditions and were maintained on AIN-93G chow (Research Diets, New Brunswick, NJ) and Pure Life water (Nestle Waters North America, Stamford, CT). Research Diets has reported levels of iAs in AIN-93G chow to be below the limit of detection, and our lab has previously confirmed levels of iAs in Pure Life water to be below the limit of detection using inductively coupled plasma mass spectrometry measurements^15,16^. Pregnant and non-pregnant female mice (6-12 weeks of age) were given Pure Life water containing 0 (control), 100 or 1000 µg/L sodium arsenite (NaAsO_2_) beginning at embryonic (E) 2.5 and continuing until harvest at E17.5 or parturition (Fig. 1). Fresh water bottles containing control or NaAsO_2_ drinking water were replenished every 2-3 days to maintain concentrations of iAs and minimize oxidation. Water and food consumption did not differ between groups (Supplemental Figure 1).

**Figure 1:**
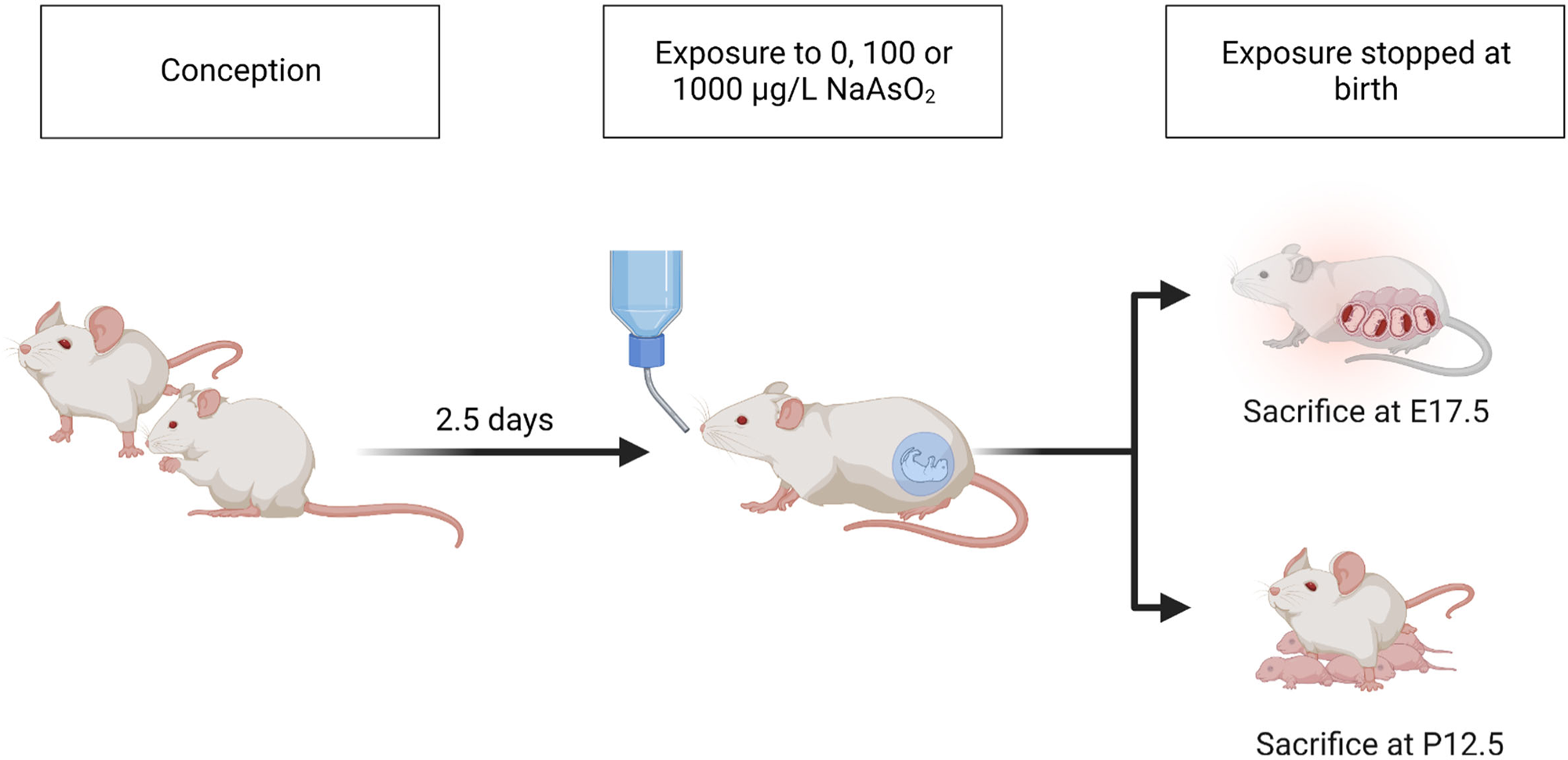
Timeline of iAs exposure. Schematic illustration of animal treatment. From embryonic day (E) 2.5 to birth, pregnant C57B6/J mice were given drinking water containing 0 (control), 100 or 1000 µg/L NaAsO_2_. At E17.5, prenatal endpoints were harvested or exposed until birth and were harvested at postnatal (P) day 12.

### Tissue Harvest and Collection

Mice were anesthetized with a mixture of ketamine (90 mg/kg; Hofspira) and xylazine (10 mg/kg; Sigma-Aldrich) via intraperitoneal injection and anticoagulated with heparin (Fresenvis Kabi, Lake Zurich, IL) prior to tissue collection. Hearts were excised, cannulated on a Langendorff apparatus, and perfused in a retrograde manner with Krebs-Henseleit buffer (95% O_2_, 5% CO_2_; pH 7.4) under constant pressure (100 cmH2O) and temperature (37°C) as previously described for 5 minutes to washout blood^40^. Buffer consisted of (in mmol/L): NaCl (120), KCl (4.7), KH_2_PO_4_ (1.2), NaHCO_3_ (25), MgSO_4_ (1.2), d-Glucose (14), and CaCl_2_ (1.75). Following perfusion, hearts were weighed and either sectioned for histology, or equally halved, and snap-frozen in liquid nitrogen and stored at −80°C for molecular studies. Tissue was collected for prepartum endpoints at E17.5, and postpartum endpoints at P12.5. Cross-fostering of offspring was not performed in this study, and dams were kept with their respective litters until P12.5.

### Echocardiography

All echocardiography procedures were performed by an investigator blinded to experimental conditions using a preclinical ultrasound imaging system (Vevo 2100; FUJIFILM VisualSonics, Toronto, ON, Canada) with a 40-MHz linear transducer, as previously described^40^. Echocardiography was performed on pregnant mice and embryonic pups under anesthesia using isoflurane. Mice were induced with 3-4% isoflurane and maintained at 1-2% thereafter. Postnatal pups underwent conscious transthoracic echocardiography. The M-mode echocardiogram was acquired from the short-axis view of the left ventricle at the level of the midpapillary muscles (200 m/s sweep speed). Semiautomated continuous tracing of the left ventricular walls from this axis view measured, calculated, or extrapolated the following cardiac parameters using 3-5 cardiac cycles: left ventricular posterior wall thickness at end diastole (LVPWd), left ventricular posterior wall thickness at end systole (LVPWs), left ventricular internal diameter at end diastole (LVIDd), left ventricular internal diameter at end systole (LVIDs), percent fractional shortening (FS), percent ejection fraction (EF), heart rate (H), and cardiac output (CO). Parameters that were calculated were done so using the LVTool function using the Vevo LAB software (VisualSonics, Fujifilm). Embryonic and maternal echocardiography was performed at E16.5 and E17.5, and at day P12 for pups.

### Histology and Cardiomyocyte Cross Sectional Area Measurement

After perfusion and gravimetric measurement, a transverse cross-section of the heart was taken including both the left and right ventricle. Tissues were fixed, embedded in paraffin, sectioned (4µM) and mounted on slides. Heart sections were stained using Hematoxylin and Eosin (H&E) to visualize cardiomyocyte integrity (2 sections) or Masson’s Trichrome to determine presence of fibrosis (4 sections), and imaged at 20x using Aperio ScanScope CS (Leica Biosystems, Deer Park, IL). Cross-sectional area of cardiomyocytes was determined manually from Masson’s Trichrome stained tissue, with 25 measurements taken from one region per transverse section across 4 sections, totaling 100 different measurements per heart. The entire transverse section was also analyzed for fibrosis expressed as a percentage of total area using an Aperio ImageScope macro (Leica Biosystems, Deer Park, IL).

### RNA Isolation, Extraction, and cDNA Conversion

Hearts were homogenized with TRIzol (1 mL, Ambion) using a bead-mill tissue homogenizer (2 × 30 s cycles, 0°C, 7,200 rpm; Precellys Evolution 24). Lysates were mixed with chloroform (200 μL, Thermo Fisher), incubated (5 min, 25°C), and centrifuged (12,000 g, 15 min, 4°C) for phase separation. RNA collected from the upper phase was precipitated by incubation (10 min, 25°C) with isopropyl alcohol (500 μL) and centrifugation (12,000 g, 8 min, 4°C). RNA pellets were washed with 75% ethanol, centrifuged (12,000 g, 10 min, 4°C), and air-dried under a laminar flow hood (30 min). Following RNA solubilization (100 μL, DEPC-treated water), RNA concentration and purity (A260/A280 range 1.98 to 2.06) were measured via spectrophotometry (1 mL sample, NanoDrop 100, Thermo Fisher). RNA was subsequently converted to cDNA per manufacturer’s instructions (High-Capacity cDNA Reverse Transcription Kit, 4368814, Thermo Fisher). Briefly, the reverse transcription reaction mix was prepared on ice, RNA (2 μg) was added, and the samples were run on a thermocycler (60 min, 37°C; 5 min, 95°C; held, 4°C; Applied Biosystems); cDNA was stored at −20°C until use.

### Quantitative RT-PCR

Expression levels of mRNA transcripts were measured using generated cDNA, a PCR (Polymerase Chain Reaction) master mix (TaqMan Fast Advanced Master Mix, Applied Biosystems) and the following validated primers (TaqMan, Applied Biosystems) on a thermocycler (2 min, 50°C; 2 min, 95°C; (1 s, 95°C; 20 s, 60°C) × 40), (QuantStudio 3, Applied Biosystems, Thermo Fisher) in a 96-well plate (USA Scientific, Beltsville MD): *Acta1* (Mm00808218_g1), *Akt1* (Mm01331626_m1), *Esr1* (Mm00433149_m1), *Esr2* (Mm00599821_m1), *Gapdh* (Mm99999915_g1), *Kncd3* (Mm01302126_m1), *Nos3* (Mm00435217_m1) *Myh6* (Mm00440359_m1), *Myh7* (Mm00600555_m1), *Nppa* (Mm01255747_g1), *Nppb* (Mm01255770_g1), *Pgrmc1* (Mm00443985_m1)*, Pgrmc2* (Mm01283154_m1)*, Prlr* (Mm04336676_m1), and *Vegfa* (Mm00437304_m1). Expression was determined using the ΔΔCT method and normalized to *Gapdh*, which did not change in cycle time with iAs treatment.

### Heart Homogenization and Protein Isolation

Hearts were homogenized in cell lysis buffer (1 mL; Cell Signaling Technology, Danvers, MA) supplemented with a protease and phosphatase-inhibitor cocktail (Cell Signaling Technology) using a hard tissue lysing kit (Precellys CK28 Lysing Kit, Bertin Instruments) with a bead-mill tissue homogenizer (2 x 30s cycles, 0°C, 7,2000 rpm; Precellys Evolution 24, Bertin Instruments). Supernatant was collected as total crude homogenate; protein concentration was determined via Bradford assay and aliquots of homogenate were stored (at −80°C until use).

### Western Blot

Protein homogenates (30 µg) were separated (20 min, 75 V; 100 min, 175 V) on a gradient Bis-Tris SDS-PAGE gel (4-12%, NuPAGE; Thermo Fisher, Carlsbad, CA) and transferred (90 min, 220 mA) to a PVDF membrane (Thermo Fisher). Every gel included two molecular-weight markers for separate regions of interest (High Range Color-Coded Prestained Protein Marker; Cell Signaling Technology; and Novex Prestained Protein Standard, Thermo Fisher) on opposite ends of the gel. Total protein served as the loading control by covalently labeling lysine residues (No-Stain Protein Labeling Reagent, Thermo Fisher) following the manufacturer’s protocol and visualized by fluorescence imaging (488 nm). Membranes were blocked (1h) with bovine serum albumin (5% wt/vol, Sigma-Aldrich) in Tris-buffered saline with Tween-20 (0.1%), and subsequently incubated (overnight, 4°C) with primary antibodies for p-Akt (S473) (1:1000, Rabbit mAb, Cell Signaling), Akt (pan) (1:1000, Rabbit mAb, Cell Signaling), p44/42 MAPK (ERK1/2) (1:1000, Rabbit mAb, Cell Signaling), and p-p44/42 MAPK (T202/Y204) (1:1000, Rabbit Ab, Cell Signaling). Membranes were then incubated with secondary antibodies (Anti-rabbit IgG, HRP-linked Ab, Thermo Fisher) and visualized using chemiluminescence substrate (SuperSignal West Pico PLUS, Thermo Fisher) and an iBright imager (Invitrogen, Thermo Fisher). Membranes were stripped with Reblot Plus Mild Solution (Millipore) when necessary to re-probe total protein following examination of phosphorylation status. No-Stain and full western blot scans are included in Supplemental Figure 2.

### Statistical Analysis

Sample sizes of mice for each experiment were estimated *a priori* via power analysis (power = 0.80, effect size = 0.25, alpha = 0.05) based on data generated in previous studies from our group. Mice were randomized during collection to minimize batch effects. All offspring data were averaged by litter to minimize litter effect. Data were analyzed using GraphPad Prism (La Jolla, CA). Statistical outliers were identified using the ROUT method (Q=1%). Statistical comparisons between groups were determined using an ordinary one-way ANOVA with Sidak’s multiple comparisons test, a two-way ANOVA with Tukey’s multiple comparisons test or a two-tailed Mann-Whitney test as appropriate. Significance was set at p < 0.05.

## RESULTS

### iAs exposure alters cardiac function in late pregnancy

To determine whether iAs exposure alters maternal cardiac structure or function during pregnancy, transthoracic echocardiography was performed at E16.5 and E17.5 during the late pregnancy period (Fig. 2). We found that pregnant dams exposed to 1000µg/L iAs had an increase in ejection fraction trending towards significance compared to pregnant control and 100 µg/L iAs exposed pregnant dams, but no changes were observed between non-pregnant exposed groups (Fig. 2A). Pregnant dams exposed to 1000 µg/L iAs also exhibited a significant increase in fractional shortening compared to control and 100 µg/L iAs exposed dams with no changes between non-pregnant exposure groups (Fig. 2B). Additionally, pregnant mice exposed to 1000 µg/L iAs showed a significant increase in cardiac output compared to control and 100 µg/L exposed pregnant dams, as well as 100 and 1000 µg/L iAs exposed non-pregnant females (Fig. 2C). Since cardiac output is the product of heart rate and stroke volume, we examined these two parameters. We found no changes in heart rate across pregnant or non-pregnant females (Fig. 2D), but stroke volume was significantly increased in 1000 µg/L iAs exposed pregnant dams compared to non-pregnant females exposed to 1000 µg/L iAs (Fig. 2E). We also found no changes in end systolic or diastolic volumes, or in average wall thickness across all pregnant and non-pregnant exposure groups (Figs. 2F-H).

**Figure 2:**
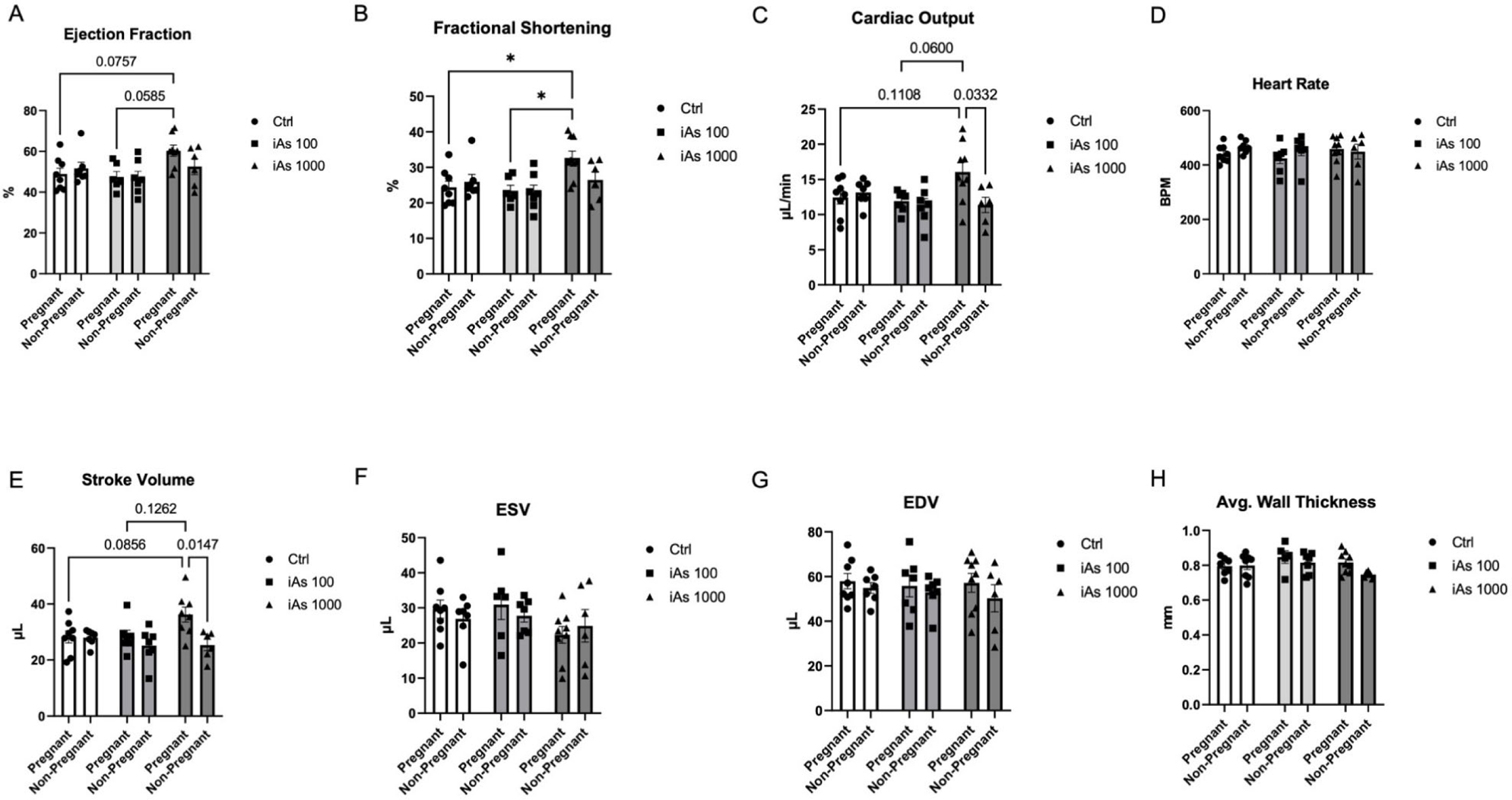
iAs exposure increases maternal cardiac function during pregnancy. Cardiac parameters as measured by transthoracic echocardiography in both pregnant and non-pregnant mice including: cardiac output (*A*), stroke volume (*B*), heart rate (*C*), ejection fraction (*D*), fractional shortening (*E*), end diastolic volume (*EDV; F*), end systolic volume (*ESV; G*), average wall thickness (*H*), left ventricular internal diameter in diastole (*LVID;d; I*), and left ventricular internal diameter in systole (*LVID;s*; *J*) (*= P < 0.05, ** = P< 0.01 vs. control; *n*=7-9 mice/group). Significance was determined by two-way ANOVA with Tukey’s multiple comparisons test.

### iAs exposure alters prepartum hypertrophic cardiac remodeling

Reversible cardiac hypertrophy is known to occur during pregnancy, so we investigated heart size across all groups to account for our observed changes in cardiac output and ejection fraction. As expected, there was a significant increase in heart size in control pregnant mice compared to control nonpregnant females (Figs. 3A, 3B). Surprisingly, during late pregnancy, we found that maternal hearts were significantly smaller in dams exposed to 1000 µg/L iAs compared to pregnant controls and were not significantly different from nonpregnant females exposed to 1000 µg/L iAs (Fig. 3A). Exposure to 100 µg/L iAs, on the other hand, was not sufficient to reduce heart growth relative to pregnant controls as heart size was increased as expected compared to non-pregnant mice with the same exposure (Fig. 3B). Next, we examined cardiomyocyte cross-sectional area in histological sections, and consistent with our observed differences in heart size, we found that cardiomyocyte cross-sectional area was significantly less in 1000 µg/L iAs-exposed hearts compared to controls (Figs. 3E, 3F). Masson’s trichrome staining was performed on transverse heart sections to examine changes in collagen deposition, but no differences were seen in collagen deposition or fibrosis (Figs. 3C, 3D).

**Figure 3:**
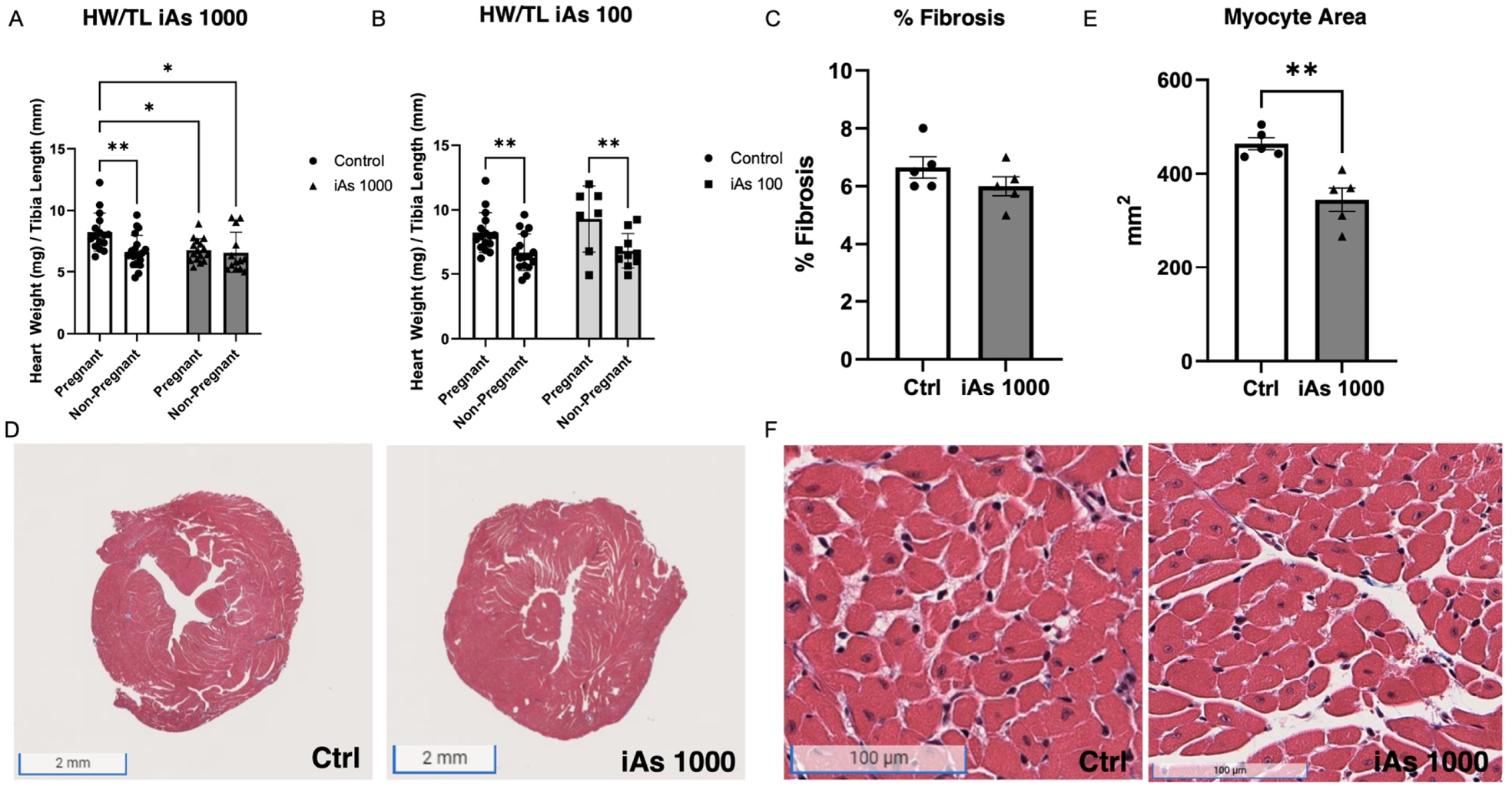
iAs exposure blunts maternal cardiac hypertrophy. Heart weight divided by tibia length in pregnant and non-pregnant females with 1000 µg/L (*A*) and 100 µg/L (*B*) iAs exposure compared to control (*n*=11-18 mice/group; *= P < 0.05, ** = P< 0.01 vs. control). Quantification of fibrosis (*C*) and representative images (*D,* scale bar 2mm) in mice exposed to 1000 µg/L iAs vs. control. Quantification of cardiomyocyte cross-sectional area (*E*) and representative images of cardiomyocyte cross-section (*F,* scale bar 100 µm) taken from 4 sections per heart with 25 cardiomyocytes per section (*n*=5 hearts/group, 100 cardiomyocytes/heart; **= P<0.01 vs. control). Significance was determined by two-way ANOVA with Sidak’s multiple comparisons test or Mann-Whitney test.

### iAs exposure during pregnancy alters the transcriptional profile of hormone receptors

Hormone signaling is important for cardiovascular adaptation and heart growth during pregnancy, so we examined the expression of various hormone receptors in the maternal heart during late pregnancy. Estrogen impacts hypertrophic remodeling in the pregnant heart^29,41^ and we found that exposure to iAs induced a dose-dependent decrease in *Esr1* (Estrogen Receptor α-ERα) mRNA levels in the whole heart (Fig. 4A). Estrogen receptor β was also probed, but the cycle time (CT) values were greater than 38, indicating low levels of expression in whole myocardium. Progesterone is also known to regulate pregnancy-induced hypertrophy, and we found that exposure to both 100 and 1000 µg/L iAs during gestation downregulated cardiac transcription of the progesterone membrane receptors *Pgrmc1* and *Pgrmc2* (Figs. 4B, 4C). Cytosolic progesterone receptors were also probed, but levels were too low in cardiac tissue to be detected. We also examined prolactin receptor expression in the heart, as prolactin is another hormone that is important during pregnancy but found that levels were not significantly altered (Fig. 4D).

**Figure 4:**
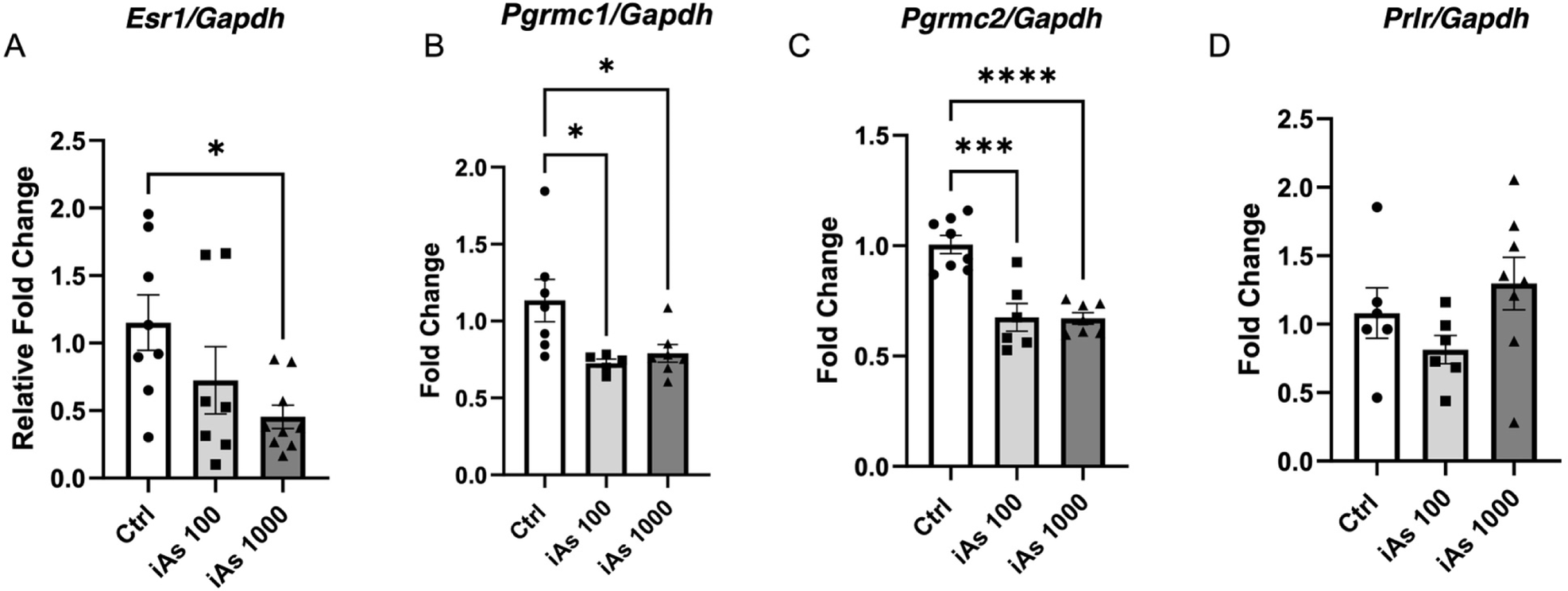
iAs downregulates estrogen receptor alpha transcripts during late pregnancy. Myocardial mRNA transcript levels of estrogen receptor alpha (*ERα*; *A*), progesterone receptor membrane component 1 (*Pgrmc1; B*), progesterone receptor membrane component 2 (*Pgrmc2; C)*, and prolactin receptor (*Prlr; D*) in pregnant females at E17.5 as measured via RT-qPCR (**= P<0.01 vs. control). Transcript levels were normalized to *Gapdh* mRNA expression, which did not change with iAs exposure. Outliers were determined by the ROUT method (Q=1%), and significance was determined by one-way ANOVA with Sidak’s multiple comparisons test or Mann-Whitney test (*n*=7-9 mice/group).

### iAs exposure and transcriptional markers of hypertrophy

Next, we examined the expression of gene transcripts typically associated with physiologic and pathologic hypertrophy, as well as genes which have been implicated in pregnancy-induced hypertrophy^42^. In hearts exposed to both 100 and 1000 µg/L iAs during pregnancy, we found a decrease in *Akt1* (Akt), a major regulator of physiologic hypertrophy that was trending towards significance (Fig. 5A). We also found a reduction in *Nppa* (ANP) at both doses (Fig. 5B). Interestingly, we found a significant reduction in *Vegfa* transcripts at 100 µg/L iAs only (Fig. 5C). Other markers of physiologic and pathologic hypertrophy were not altered (Fig. 5D-I). Transcriptional results were confirmed via western blot and protein levels of Akt were significantly reduced in 1000 µg/L iAs exposed hearts, while Akt phosphorylation (Ser473) relative to total Akt showed a trending decrease (Fig. 6A and 6B). Extracellular-regulated kinase 1 and 2 (ERK1/2) is also a major regulator of pregnancy-induced hypertrophy^43^. However, we found that levels of ERK1/2, as well as phosphorylation of the activation site, were unchanged with iAs exposure (Fig. 6C and 6D).

**Figure 5:**
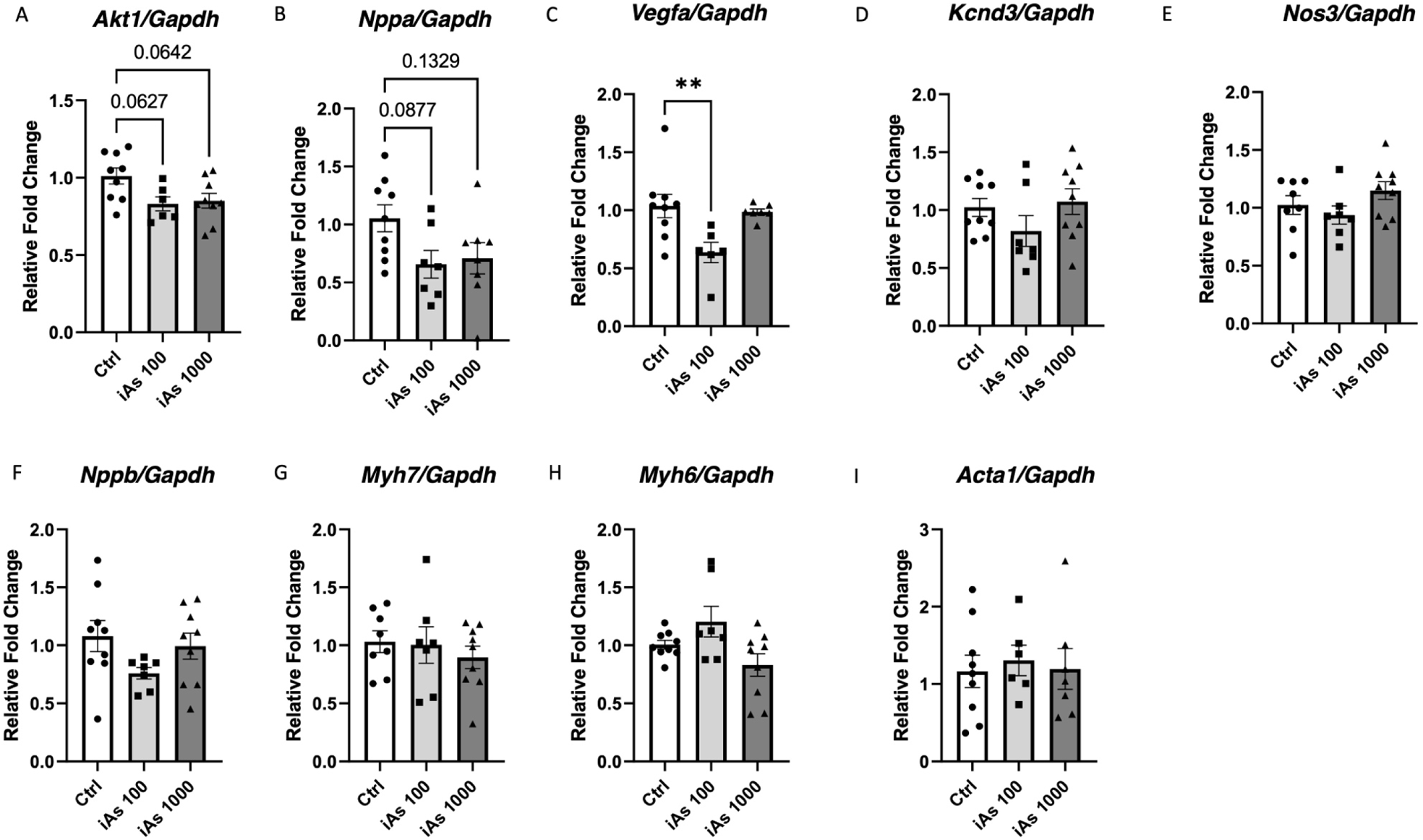
iAs exposure reduces Akt and ANP transcripts in the prepartum heart. Myocardial mRNA transcript levels of protein kinase B (*Akt; A*), natriuretic peptide A (*ANP; B*), vascular endothelial growth factor (*Vegf; C*), potassium voltage-gated channel 4.3 (*Kv4.3; D*), endothelial nitric oxide synthase (*Nos3; E*), natriuretic peptide B (*BNP; F*), β-myosin heavy chain (*Myh7; G*), α-myosin heavy chain (*Myh6; H*), skeletal muscle α-actin (*Acta1; I*), (*=P<0.05, *n*=7-9 hearts/group). Outliers were determined by the ROUT method (Q=1%), and significance was determined by a one-way ANOVA with Sidak’s multiple comparisons test.

**Figure 6:**
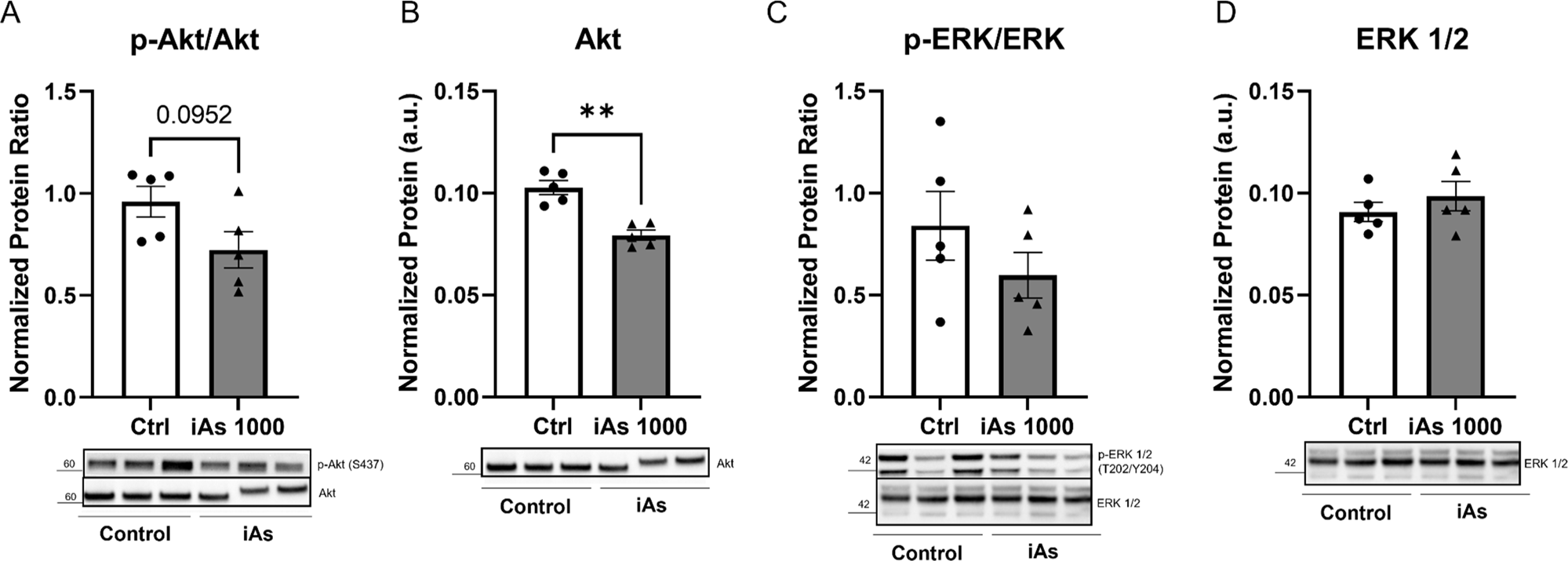
iAs exposure reduces protein levels of Akt in the prepartum heart. Myocardial protein expression of phosphorylated Akt at the activating site, S437, normalized to total Akt (*A*) in dam hearts (P= 0.0776 vs. control, *n=* 5 hearts/group). Protein expression of total Akt (*B*) in dam hearts (***P=0.0005 vs. control). Protein expression of phosphorylated ERK1/2 at the activating sites, T202/Y204 (*C*), and of total ERK1/2 (*D*). Protein levels of each target were normalized to total transferred protein levels. Outliers were determined by the ROUT method (Q=1%), and significance was determined by a Mann-Whitney test.

### In utero exposure to iAs and offspring outcomes

Finally, we examined the impact of gestational iAs on offspring outcomes with a focus on cardiac development. We found that in utero exposure to 100 or 1000 µg/L iAs during gestation resulted in no change to litter size (Fig. 7A) or embryonic body weight (Fig. 7B). However, heart weight relative to body weight was increased at E17.5 after exposure to 1000 µg/L iAs (Fig. 7C). To further delineate changes in cardiac structure and function, echocardiography was performed in utero at E17.5, but we found no changes in average wall thickness (Fig. 7D), LVPWd (Fig. 7E), IVSd (Fig. 7F), EF (Fig. 7H), or FS (Fig. 7I). However, we did find an increase in the left ventricular internal diameter during diastole at E17.5 with exposure to 1000 µg/L IAs (LVIDd Fig. 7G).

**Figure 7:**
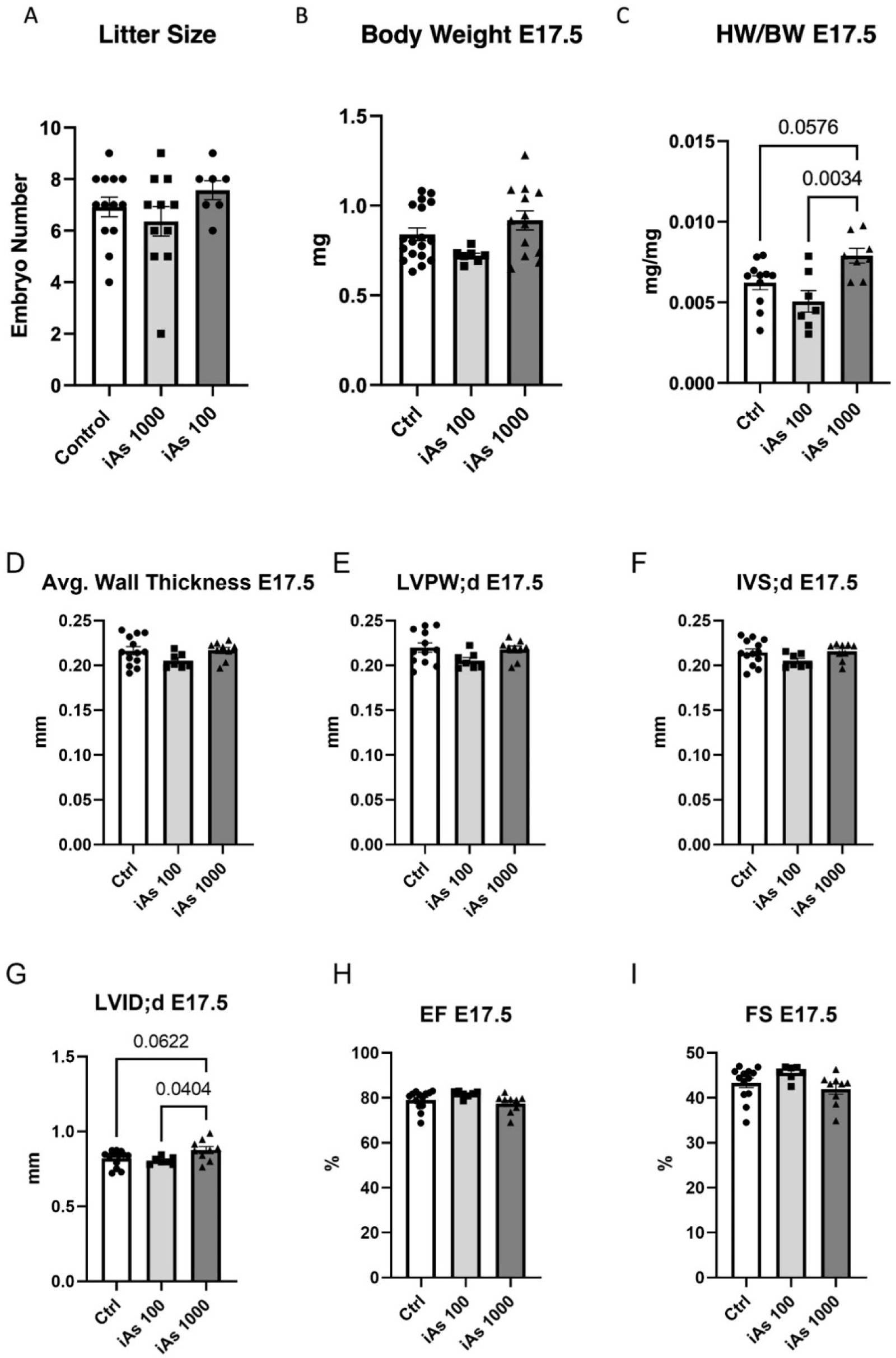
Offspring from iAs exposed dams show increased heart weight and left ventricular internal diameter at E17.5. Litter size (*A*), body weight (B), and embryonic heart weight normalized to body weight (*C*), in offspring exposed to 100 and 1000 µg/L iAs during gestation (*n*= 16-18 litters/group). Cardiac parameters measured in-utero using echocardiography including average wall thickness (D), left ventricular posterior wall thickness (LVPW; E), interventricular septum thickness (IVS; F), left ventricular internal diameter (LVID; G), ejection fraction (EF; H), and fractional shortening (FS; I). Outliers were determined by the ROUT method (Q=1%), and significance was determined by one-way ANOVA with Sidak’s multiple comparisons test.

These same parameters were measured again postnatally at P12, and we found that exposure to 100 µg/L iAs in-utero reduced body weight at P12 (Fig. 8A). When stratified by sex, both sexes showed a similar significant reduction in growth (Fig. 8B). We also saw no differences in distribution of sex by exposure (Supplemental Table 1). However, while an increase in heart weight was noted at E17.5, we found no changes compared to controls at P12, with or without sex stratification (Figs. 8C, 8D). Echocardiography also revealed no changes in average wall thickness (Fig. 8E), IVS (Fig. 8F), LVPW (Fig. 8G), LVIDd (Fig. 8H), EF (Fig. 8I), or FS (Fig. 8J).

**Figure 8:**
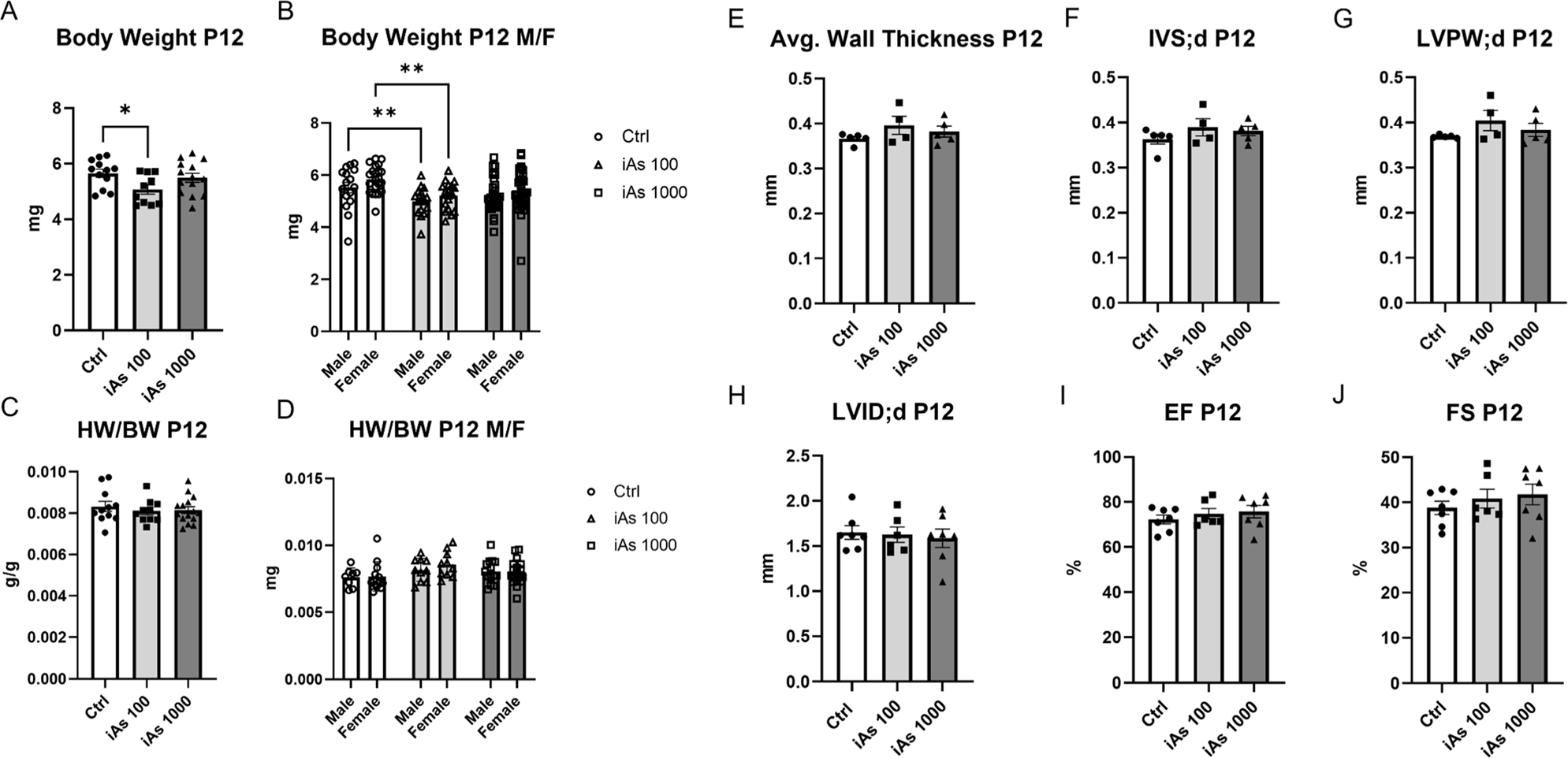
Offspring exposed to iAs in-utero have reduced body weight at P12. Body weight at P12 (A), body weight stratified by sex (B), heart weight relative to body weight (C), and heart weight relative to body weight stratified by sex (D) measured gravimetrically. Cardiac function parameters measured via echocardiography including average wall thickness (E), interventricular septum thickness (IVS; F), left ventricular posterior wall thickness (LVPW; G), left ventricular internal diameter (LVID; H), ejection fraction (EF; I), and fractional shortening (FS; J). Significance was determined by one-way ANOVA with Sidak’s multiple comparisons test.

## DISCUSSION

We report here for the first time that iAs exposure during gestation at environmentally relevant doses blunts pregnancy-induced hypertrophy in the maternal heart during late pregnancy and leads to the downregulation of hormone receptors ERα and PGRMC1/2, as well as ANP and Akt expression. Pregnancy-induced cardiac hypertrophy is critical for meeting the circulatory demands of the mother and the developing fetus during gestation, and we find that iAs exposure prevents this important hypertrophic process from occurring properly^18,44^. We observed a compensatory change in cardiac function in iAs-exposed dams (Fig. 2), despite the lack of proper heart enlargement (Fig. 3), along with downregulation of ERα, PGRMC1/2, (Fig. 4) and ANP and Akt (Figs. 5 and 6). Additionally, we found an increase in heart weight relative to body weight in offspring (Fig. 7) and decreased growth at P12 (Fig. 8) suggesting that either direct iAs exposure or improper maternal cardiac remodeling may contribute to adverse outcomes in offspring.

While our previous study demonstrated that iAs induced sex-dependent changes in cardiac hypertrophy, these changes were exclusive to adult males and adult females remain unaffected after an 8-week exposure^16^. However, in the current study, we find that pregnant females are especially vulnerable to alterations in cardiac hypertrophy with gestational iAs exposure. This study is the first to examine the impact of iAs exposure on maternal cardiac remodeling during pregnancy and our findings identify a critical window of exposure that leads to a dysregulation in pregnancy-induced maternal cardiac growth and endocrine signaling with iAs exposure.

### iAs exposure and maternal cardiac function

Heart growth or hypertrophy is a critical process that occurs during pregnancy. Prior work has demonstrated an increase in heart weight and cardiomyocyte cross-sectional area in mice during late pregnancy^45^. Consistent with these findings, we found that control or 100 µg/L iAs-exposed dams showed a significant increase in heart weight compared to iAs-exposed and non-exposed non-pregnant females (Fig. 3). However, cardiac growth was blunted in maternal hearts exposed to 1000 µg/L iAs during pregnancy, as we noted significantly smaller heart weight with 1000 µg/L iAs exposure compared to control and 100 µg/L iAs exposed dams (Fig. 3). Cross-sectional area was also significantly lower in cardiomyocytes from 1000 µg/L iAs exposed dams compared to non-exposed pregnant controls (Fig. 3), providing further confirmation that pregnant dams exposed to 1000 µg/L iAs during gestation have smaller hearts during late pregnancy. Left ventricular hypertrophy during pregnancy results from volume overload, and a lack of hypertrophy vs. pregnant controls may indicate a lack of blood volume expansion^44,46^. As such, plasma volume will be examined in future studies at different timepoints during pregnancy.

Furthermore, we found that cardiac function was enhanced in pregnant dams exposed to 1000 µg/L iAs compared to pregnant controls or dams exposed to 100 µg/L iAs (Fig. 2), likely as a compensatory mechanism to increase blood flow to the mother and developing fetus. This increase in cardiac function in 1000 µg/L iAs-exposed dams may result from a number of mechanisms that include enhanced cardiomyocyte contractility or increased beta-adrenergic stimulation and future studies will examine these potential compensatory mechanisms. Importantly, sustained alterations to either of these pathways could result in detrimental changes to the heart which could include a transition from pregnancy-induced hypertrophy to pathological hypertrophy or increased risk for prolonged cardiovascular disease postpartum^47^. As such, postpartum remodeling and specifically the impact of iAs exposure during pregnancy on the postpartum heart, is a critical area that needs to be addressed in future mechanistic studies. Importantly, we find no alterations in cardiac size or function with iAs exposure alone in non-pregnant females at E17.5, or 15 days of exposure (Figs. 2 and 3). We also found no change to cardiac structure and function in adult female mice in our prior studies examining iAs exposure at 4 and 8 weeks^15,16^. Taken together, our findings suggest that the unique combination of iAs exposure and pregnancy are necessary for increasing cardiac function (Fig. 2) and blunting pregnancy-induced cardiac growth (Fig. 3).

### iAs exposure decreases myocardial hormone receptor expression

Pregnancy-induced hypertrophy is governed largely by sex-hormones, such as progesterone and estradiol, and previous research has shown that iAs can act as an endocrine disruptor^48–51^. In the current study we found that exposure to 1000 µg/L iAs downregulated cardiac ERα, Pgrmc1, and Pgrmc2 during late pregnancy (Fig. 4). Prior work has shown that iAs exposure downregulates various steroid hormone receptors and decreases ERα expression in the uterus, leading to alterations in uterine hypertrophy^49,52^. Estradiol has been shown to be anti-hypertrophic in mice at high levels and pro-hypertrophic at low levels in cultured cells, mediating the activation of certain downstream signaling pathways that are crucial for cardiac hypertrophy^25,42,43^. Exposure to iAs has also been shown to impact progesterone signaling, decreasing serum progesterone in rats as well as progesterone receptors in culture^53–55^. Consistent with these findings, our data showed decreased transcripts of receptors for progesterone and estradiol in the heart (Figs. 4A, 4B, 4C). Interestingly, we observed that doses of 100 and 1000 µg/L iAs downregulated Pgrmc1 and Pgrmc2 at equal levels, while a dose-dependent reduction in ERα was noted with 100 and 1000 µg/L iAs. Since we only observed maladaptive changes to physiologic hypertrophy at 1000 µg/L iAs, altered ERα transcription, either alone or in combination with the downregulation of Pgrmc1 and Pgrmc2 transcripts, likely contributes to the lack of cardiac hypertrophy observed in the maternal heart.

Dampened estradiol signaling during pregnancy can impact blood volume expansion and the ability of the heart to grow in size. Cardiac hypertrophy during pregnancy is reliant upon major signaling pathways which are activated by a combination of mechanotransduction due to increased blood volume and hormone signaling^25,43,56^. Plasma volume increase is a consequence of sodium resorption and water retention, signaled through placental estrogen production^57^. The ability of iAs to act as an endocrine disruptor and downregulate ERα expression in the maternal heart during pregnancy may explain why we only see increased cardiac function and reduced heart size in pregnant mice exposed to 1000 µg/L iAs. However, further studies are necessary to characterize the impact of iAs on hormonal changes at different time points throughout pregnancy.

### Akt and ANP reduction in maternal hearts exposed to IAs

Transcriptionally, pregnancy-induced hypertrophy has been found to resemble exercise-induced hypertrophy, but there are some notable differences^24^. For example, genes such as Acta1, ANP, and BNP are found to be upregulated in late pregnancy ^24,58,59^. When probing for genes associated with pregnancy-induced cardiac hypertrophy^42,44^ we found that dams exposed to either 100 or 1000 µg/L iAs showed reduced maternal heart expression of Akt (Fig. 5A), a primary mediator of pregnancy-induced hypertrophy^43^. We also found that protein expression of Akt was reduced in the maternal heart, and phosphorylation of the active site showed a trending decrease with iAs exposure (Figs. 6A, 6B). Interestingly, we observed an equal reduction in the transcription of Akt at both doses, but we only observed phenotypic changes in dams exposed to 1000 µg/L iAs. This may be due to alterations in other molecular pathways at the 1000 µg/L iAs dose (i.e., hormone signaling). Additional studies have shown that exposure to iAs in adult rats reduces Akt expression in cardiac tissue irrespective of pregnancy^60^. While iAs exposure reduces myocardial Akt expression in our model, additional research is needed to determine how this reduction in Akt expression specifically contributes to the blunted hypertrophy observed in iAs-exposed pregnant mice.

Exposure to 100 or 1000 µg/L iAs also induced a ∼50% reduction in transcript levels of ANP in the maternal heart (Fig. 5B). ANP is secreted by the heart in response to atrial distension and is found to be increased consistently in blood plasma throughout pregnancy^58,61,62^. ANP is a vasodilator and transcription is induced by atrial stretching due to volume overload^63^. These findings further suggest that iAs exposure could alter plasma volume expansion during pregnancy. In humans, exposure to iAs increased hemoglobin concentration and decreased blood protein concentration, indicating that iAs exposure may affect plasma volume regulation^64,65^. However, this has not been explored in the context of pregnancy. While a reduction in both ANP and Akt are associated with cardiovascular alterations during pregnancy, the reduction in transcription at both doses and the resulting pathophysiologic changes only at the highest dose may have several implications. It is possible that ANP and Akt are acting in a synergistic manner with our observed changes in hormone receptor signaling. The other possibility is that alterations to ANP and Akt levels at this timepoint are not associated with the changes in hypertrophic signaling. However, the full molecular pathway has yet to be elucidated, and understanding how the transcriptional and expression profile of the maternal heart changes with iAs exposure during early- and mid-pregnancy will be essential to further our understanding of the full impact of iAs.

### Effects of gestational IAs exposure on offspring

In the offspring we found that heart weight was increased with 1000 µg/L iAs exposure, but not with 100 µg/L iAs (Fig. 7C). Left ventricular diameter was also increased at E17.5 with 1000 µg/L iAs exposure (Fig. 7G). In mammals, heart cells proliferate from conception until late gestation and switch to cell hypertrophy shortly after birth^66^. Pregnancies with metabolic dysfunction, such as obesity and diabetes, result in increased fetal cardiac hypertrophy resulting from increased cardiomyocyte cell size^67–69^. In adult hearts, enlarged left ventricular diameter in combination with decreased ejection fraction culminates in heart failure^70^. However, we do not see changes to ejection fraction with either iAs exposure dose (Fig. 7H), which suggests that there are only structural changes to the heart at E17.5. Several epidemiologic studies have associated in-utero exposure to iAs with congenital heart defect^71–73^, but this has yet to be replicated in a murine model with a relevant exposure dose or route of exposure. We also examined postnatal cardiac function at P12 but found no alterations to cardiac function or structure across doses (Fig. 8). These findings suggest that changes observed to the fetal heart at E17.5 may prime the heart for disease as an adult. Many epidemiologic studies have found that in-utero exposure to iAs induces cardiovascular disease later in life^74–77^.

Furthermore, we did not see alterations to litter size, embryonic body weight, or differences in sex distribution of litters at E17.5 with exposure to either 100 or 1000 µg/L iAs beginning at E2.5 (Fig. 7A, 7C). In humans, in-utero exposure to iAs has been shown to increase spontaneous abortion and decrease birthweight^78–80^. While it is still unclear if this phenotype is seen in mice, it is possible that disrupted embryonic growth is due to exposure occurring before E2.5. We characterized offspring body and organ growth postnatally at P12 and found that exposure to 100 µg/L iAs in-utero impairs growth as measured by body weight at P12 (Fig. 8A). Similar findings were observed in a prior study using doses as low as 10 µg/L iAs ^81^. However, that study showed that cross-fostering the pups reduced the growth deficit, suggesting that the reduction in growth was due to breast milk nutrition rather than altered in-utero programming^81^. Since our model halted iAs exposure at birth, and we did not cross-foster, it is possible that iAs may have lasting effects on breast milk nutrition. Additionally, iAs has been shown to alter epigenetic re-programming in offspring after in-utero exposure^82,83^. In human studies, iAs has been shown to regulate birthweight via placenta-derived miRNAs^84,85^. While we see evidence of some phenotype in offspring exposed to iAs in-utero, more experiments are needed to define exposure windows and understand how these changes persist into adulthood and underlying signaling mechanisms.

## LIMITATIONS

While this study provides important insights into the effects of iAs on maternal health, there are limitations that require acknowledgement. Firstly, this study uses only a single prepartum timepoint, so future studies will characterize physiology during early and mid-pregnancy timepoints throughout the 21-day window of murine pregnancy. Exposure was started at E2.5 during gestation, while a real-world exposure would encompass a lifetime. Although this is a limitation of our study, this exposure scenario is important for defining a critical window of maternal exposure. Additionally, experiments were conducted using a mouse model, and while mice are well characterized and commonly utilized for studies examining pregnancy and iAs exposure, it is important to note that iAs metabolism in a mouse is different in mice compared to humans^86–88^. The mouse metabolizes iAs more quickly than humans, making higher doses in mice equivalent to lower doses in human populations. There are also key differences in murine pregnancy compared to that of humans. Many rodent studies have not been able to replicate some of the findings associated with in-utero exposure of iAs on development in humans, indicating that there are differences between the two species, and this should be taken into consideration when interpreting these findings. iAs has also been shown to target mitochondria and affect cellular metabolism^89–94^. Shifts in metabolism in the heart during pregnancy have been shown to be essential for cardiac hypertrophy^45,95–99^. While we did not examine iAs-dependent effects on myocardial metabolism in the current study, these pathways need to be further investigated. Finally, we only examined two doses in our study, 100 µg/L and 1000 µg/L iAs. Lower doses should be used in future studies to understand if these effects, specifically postpartum, persist at lower doses, such as the recommended limit of 10 µg/L provided by the Environmental Protection Agency.

## CONCLUSION

iAs exposure during gestation blunted pregnancy-induced hypertrophy and increased cardiac function during late pregnancy, likely via downregulation of ERα, PRMGC1/2, ANP and Akt. As such, these changes have the potential to contribute to adverse pregnancy-related cardiovascular events, impact the developing offspring, as noted, and may lead to increased risk for maternal CVD postpartum. Taken together, our findings suggest that exposure to iAs during pregnancy causes adverse effects on both the pregnant dam and the offspring. Our findings also add to the growing body of evidence implicating iAs as a driver of cardiovascular disease and a potential risk factor for maternal and fetal health during pregnancy, underscoring the importance of removing iAs from human consumption.

## ACKNOWLEDGEMENTS

We acknowledge the technical assistance of the Oncology Tissue Services Core at the Johns Hopkins University School of Medicine and the Small Animal Cardiovascular Phenotyping and Model Core at the Johns Hopkins University School of Medicine.

## FUNDING

This work was supported by the National Institutes of Health [T32 ES007141 (NT), F31 HL165820 (HG), R21 HL157800 (MK) and R01 HL136496 (MK)].

## Notes

### Competing Interest Statement

The authors have declared no competing interest.

